# Decoding Hierarchical Control of Sequential Behavior in Oscillatory EEG Activity

**DOI:** 10.1101/344135

**Authors:** Atsushi Kikumoto, Ulrich Mayr

## Abstract

Despite strong theoretical reasons for assuming that abstract representations organize complex action sequences in terms of subplans (chunks) and sequential positions, we lack methods to directly track such content-independent, hierarchical representations in humans. We applied time-resolved, multivariate decoding analysis to the pattern of rhythmic EEG activity that was registered while participants planned and executed individual elements from pre-learned, structured sequences. Across three experiments, the theta and alpha-band activity independently coded basic elements *and* abstract control representations, in particular the ordinal position of basic elements, but also the identity and position of chunks. Further, a robust representation of higher-level, chunk identity information was only found in individuals with above-median working memory capacity, potentially providing a neural-level explanation for working-memory differences in sequential performance. Our results suggest that by decoding oscillations we can track how the cognitive system traverses through the states of a hierarchical control structure.

---

Whether we dance or compose music, write computer code, or plan a speech: A limited number of basic elements––dance moves, notes, code segments, or arguments––need to be combined in a goal-appropriate order. Following Lashley (1951), many theorists believe that such sequential skills are achieved through a set of abstract, content-independent control representations that code the position of basic elements within shorter subsequences (i.e., chunks) or the position of chunks within a hierarchically organized plan (Dehaene, Meyniel, Wacongne, Wang, & Pallier, 2015; Rosenbaum, Kenny, & Derr, 1983).

Hierarchical control representations have been proposed as the hallmark of complex, flexible behavior in humans, including motor or action control (Collard & Povel, 1982; Cooper & Shallice, 2006; Rosenbaum et al., 1983; Schneider & Logan, 2006), language (Fitch & Martins, 2014), or problem solving (Carpenter, Just, & Shell, 1990; Miller, Galanter, & Pribram, 1986). However, in principle, serial order can also be established through more parsimonious mechanisms based on “associative chaining” between consecutive elements, without having to assume content-independent representations (Botvinick & Plaut, 2004; Davachi & DuBrow, 2015). Empirically, it is exactly the abstract nature of serial-order control representations that has made it difficult to provide direct evidence for their existence, to distinguish them from chaining-based representations, or to characterize their functional properties.

The current work tests the general hypothesis that the pattern of frequency-band specific rhythmic activity in the EEG reflects different types of serial-order control representations and allows tracking the current position within a hierarchical control structure. This hypothesis is based on two sets of observations.

First, hierarchical control representations need to coordinate a large array of sensory, motor, and higher-level neural processes. A number of results suggest that such dynamic, and large-scale neural communication can be achieved through frequency-specific, rhythmic activity that coordinates local and global cellular assemblies (Buzsaki, 2006; Fries, 2005; Helfrich & Knight, 2016). For example, the power of cortical oscillations in the theta band (4 to 7 *Hz*) increases as a function of serial-order processing demands (Gevins, Smith, McEvoy, & Yu, 1997; Jensen & Lisman, 2005; Jensen & Tesche, 2002; Roberts, Hsieh, & Ranganath, 2013; Sauseng et al., 2009). Specifically, Hsieh, Ekstrom, and Ranganath (2011) reported that short-term memory of the order of serially presented visual objects was associated with increased frontal theta-band power. In contrast, the memory of the items themselves was related to changes in posterior alpha power (8 to 12 *Hz*). Combined, these results indicate that both the content and the serial order of a sequence are represented in oscillatory activity. Furthermore, activity in the alpha-band may code for basic elements and in the theta-band for how elements are ordered.

Second, goal-relevant properties induce large scale neural responses that are specific to the contents of representations (Barak, Tsodyks, & Romo, 2010; Stokes, Wolff, & Spaak, 2015). These neural response profiles in turn can be decoded with time-resolved, multivariate pattern analysis (MVPA) from the pattern of spectral-temporal profile of EEG or MEG activity (Foster, Sutterer, Serences, Vogel, & Awh, 2016; Fukuda, Kang, & Woodman, 2016). While the majority of this research has focused on the representation of basic perceptual properties, there is also some evidence that more abstract information, such as decision confidence (King, Pescetelli, & Dehaene, 2016) or semantic categories (Chan, Halgren, Marinkovic, & Cash, 2011) can be decoded from the pattern of activity distributed over the scalp. We test here the hypothesis that even abstract, serial-order control representations can be decoded from the EEG signal independently of the basic elements.

By tracking representational dynamics while sequential action unfolds, we can also address open questions about the architecture of serial order control. Traditional models of hierarchical control are informed by the characteristic response time (RT) pattern during sequence production, with long RTs at chunk transitions and bursts of fast, within-chunk responses (Collard & Povel, 1982; Rosenbaum et al., 1983). This pattern may indicate that higher-level, chunk representations are needed only during chunk transitions, after which control is handed down to representations that code for the position and elements within chunks. Alternatively, higher-level codes may need to remain active while lower-level representations are being used—in order to ward off interference from competing chunks, or to maintain the ability to navigate within the overall control structure once within-chunk processing has completed (Ranti, Chatham, & Badre, 2015). Currently, no neural-level data are available to distinguish between these theoretical options.

A related goal is to identify the source of individual differences in sequential performance within the serial-order control architecture. Initial evidence points to the importance of working memory (WM) capacity as one limiting factor during sequential performance, at least for longer sequences (Bo & Seidler, 2009). However, little is known about the exact nature of these constraints. One possibility is that WM limits the quality of representations in a uniform manner. This would imply that decodability of all representations, no matter which type (i.e., content or structural), or on which level, is reduced for individuals with low WM capacity. Arguably however, the chunking of larger sequences into smaller subunits occurs precisely to protect representations from WM-related constraints (Brady, Konkle, & Alvarez, 2009; Cowan, 1988; Oberauer, 2009). Thus, based on this view, individual differences in sequential performance should be determined by the degree to which higher-level representations can be maintained, while action selection is controlled by within-chunk representations. This would lead to the prediction that low WM capacity selectively affects decodability of higher-level representations of chunk identity and/or position, but does not affect lower-level, within-chunk representations.

In humans, there is already some evidence from studies using short-term or recognition memory tasks and fMRI neuroimaging for ordinal position codes (Desrochers, Chatham, & Badre, 2015; Heusser, Poeppel, Ezzyat, & Davachi, 2016; Kalm & Norris, 2014; Lehn et al., 2009). However, as recently argued by Kalm and Norris (2017a), caution is in order when interpreting these results. Specifically, with the combination of short-term memory tasks and the low temporal resolution of fMRI, it is difficult to rule out serial-position confounds, such as sensory adaptation, temporal distance from a context change (i.e., beginning of a probe period), or differences in processing load (i.e., higher retrieval demands or uncertainty at the start of a sequence or chunk). Further, even if position codes can be validly decoded from the fMRI signals, it is difficult to draw strong conclusions about the temporal dynamics between different serial-order control representations. Similarly, while fMRI neuroimaging studies have yielded important insights about the neural basis of different levels within a hierarchical control structure (Badre & D’Esposito, 2009; Farooqui, Mitchell, Thompson, & Duncan, 2012; Koechlin & Summerfield, 2007), we know very little about when in the course of sequential performance which control representations are actually used.

With the analysis of EEG signals, we are in a better position to track the temporal dynamics of serial-order control codes than when relying on fMRI analyses. In addition, as experimental task, we use a procedure in which participants were asked to repeatedly cycle through an explicitly instructed sequence of simple mental operations. Previous behavioral work has provided strong behavioral evidence for the use of position codes within this paradigm (Mayr, 2009). Importantly, the cycling-sequence procedure prevents some of the most important position confounds, such as sensory adaptation or temporal distance from context changes. By allowing tight temporal control over when, which operations need to be carried out, it also provides means for dealing with the most important processing-load confounds identified by Kalm and Norris (2017a).

## Results

### Overview

We measured EEG while participants performed an explicit sequencing task that was modeled after the task-span paradigm (Mayr, 2009; Schneider & Logan, 2006). For each block of trials, participants were instructed to “cycle through” a sequence of nine orientation judgements, where the specific, expected target orientations served as basic sequential elements. Sequences had a 2-level, hierarchical structure. Level 1 consisted of three possible chunks of three ordered elements (45°, 90°, 135° orientations). On Level 2, we used either three (Experiments 1 and 2) or two (Experiment 3) ordered chunks within an overall, repeating sequence (Figure 1a and 1b). Thus, assuming the existence of Lashley-type control codes, we should expect on Level 1, independent representations that code for the basic sequential elements (i.e., orientations) and for the position of each element within a chunk. On Level 2, we attempted to identify representations that code for the identity of each chunk and of also for its position within the larger sequence.

**Figure 1.**
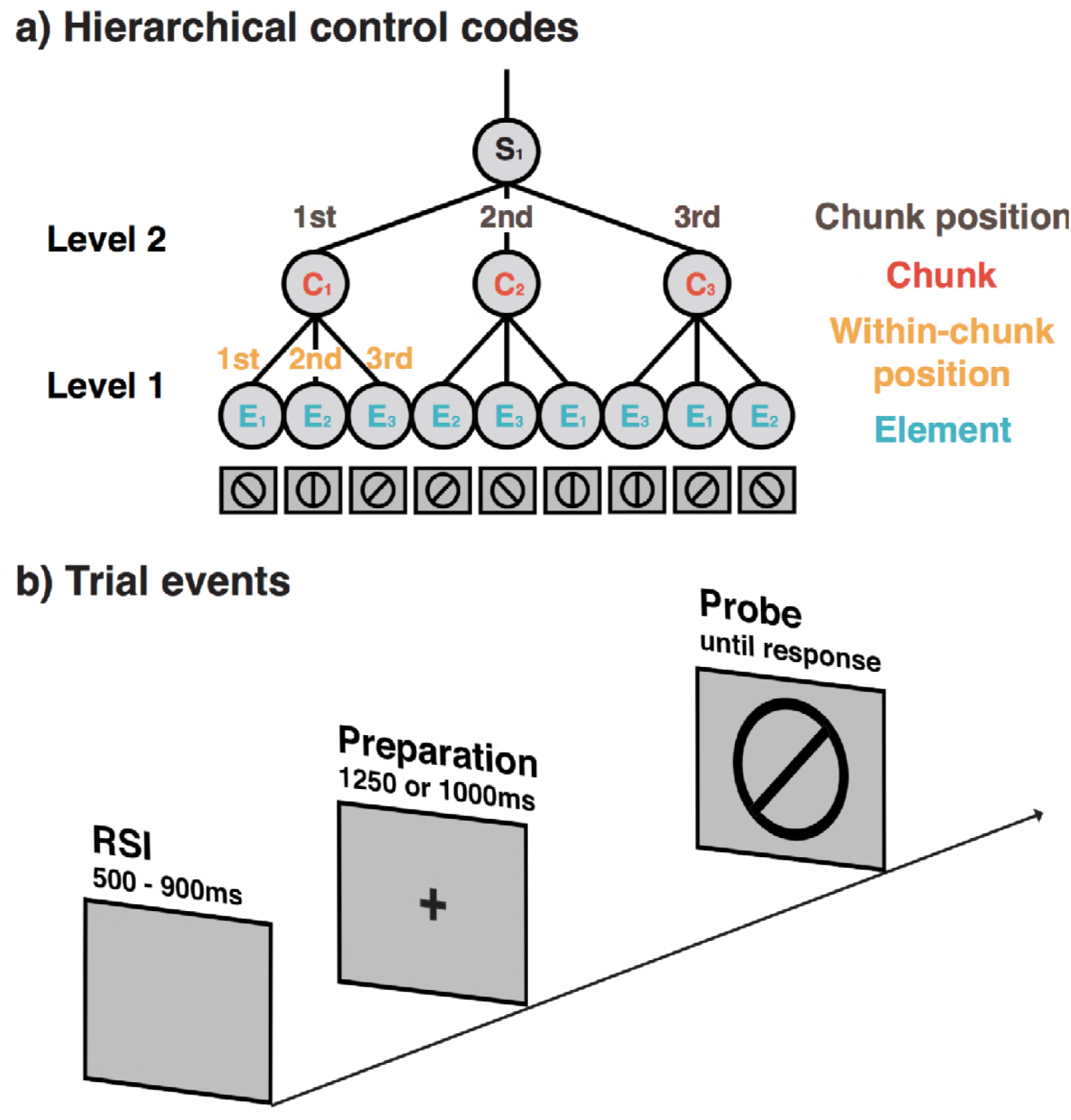
(a) Example of a 9-element sequence as used in Experiments 1 and 2, including the hierarchical structure, and the terminology for different representations used throughout. (b) Structure of individual trials in Experiments 1 and 3. Trials in Experiment 2 included a self-paced retrieval period prior to the preparation period.

As mentioned in the Introduction, identifying neural indicators of sequential representations, in particular of position codes, is particularly challenging because there may be confounding variables that differ across positions within a sequence (Kalm & Norris, 2017a). The cycling-sequence paradigm eliminates some of the position confounds, such as sensory adaptation or the temporal distance from an encoding or retrieval phase. However, another set of confounds arises because early positions within a chunk are likely to invoke higher retrieval or WM demands. To counter such processing-demand confounds, we used in Experiment 2 a self-paced procedure, where participants were asked to move per key press on to the next position in the sequence, only after they retrieved the upcoming element. The rationale here is that the self-paced retrieval time should absorb processing demand effects and remove them from the actual preparation period (during which EGG registration occurred). In Experiment 3, we returned to a fixed-paced procedure, but reduced processing-demand effects by providing more robust pretraining of each chunk, prior to the actual test block.

In Experiments 1 and 2, all sequences were constructed from just three different chunks. The use of three chunks was motivated by the goal of establishing a level playing field for decoding all different types of codes (i.e., three elements, three within-chunk positions, three chunks, and three chunk positions). However, the use of only three chunks and the constraint of counterbalancing individual elements (i.e., orientations) across positions, resulted in within-chunk sequences that contained the same element-to-element transitions across chunks (just at different positions; e.g., ***AB***C and C***AB***). In principle, the representation of such sequences could be supported through a simple chaining mechanism that requires no abstract position codes. In order to generalize our position decoding results to a situation where chaining is not possible, we used in Experiment 3 six possible sequences. Here, participants experienced each element-to-element transition equally often (i.e., ABC, ACB, BAC, BCA, CAB, ABA), rendering a simple chaining mechanism that picks up on across-sequence regularities useless.

In order to identify the representations that control sequential performance, we performed a series of decoding analyses with the spectral-temporal profiles of EEG activity across the scalp. We examined the entire preparation period prior to the appearance of the test probe (i.e., 1250 ms for Experiment 1, 1100 ms for Experiment 2) and the initial 300 ms following the test probe. At each time sample during these periods, the pattern of rhythmic activity at a specific frequency range (from 4 Hz to 35 Hz) was used separately to train linear classifiers via a 4-fold cross-validation procedure. This generated a matrix of decoding results (i.e., frequency by time samples) for each of the task-relevant representations:(1) elements, (2) within-chunk ordinal positions, (3) chunk identities, and (4) chunk positions (Figure 1a).

### Experiments 1 and 2

Experiments 1 and 2 used very similar design and procedures and therefore will be reported together. RTs from error-trials, post-error trials, and trials in which RTs were longer than 100 ms were excluded from analyses. For Experiment 2, trials with a self-paced retrieval time longer than 8000 ms were excluded. To analyze the effects of ordinal positions on serial-control performance, we specified two sets of orthogonal contrasts for positions within chunks, the first comparing position 1 against positions 2 and 3, the second comparing positions 2 and 3.

### Behavior

We examined the pattern of RTs and errors in order to confirm that participants used a hierarchical structure to perform the sequencing task (Figure 2). Specifically, we examined to what degree participants were slower and less accurate at the first position of a chunk, compared to the remaining positions. Note, that for both experiments, but in particular for Experiment 2 we had emphasized to participants the need to prepare for each upcoming probe. Thus, compared to previous work with similar sequencing tasks (e.g., Mayr, 2009) we had expected rather subtle structure effects in RTs, whereas we expected errors to remain sensitive to the greater difficulty of activating the correct chunk during chunk boundaries. For Experiment 1, which used the fixed-paced procedure, RTs showed a small, but reliable chunk boundary effect of 14 ms (*SD*=18) in RTs, *F*(1,28)=19.20, *MSE*=622.47, *p*<.001, and a robust error effect of 3.8% (*SD*=4.3), *F*(1,28)=22.70, *MSE*<0.01, *p*<.001.

**Figure 2.**
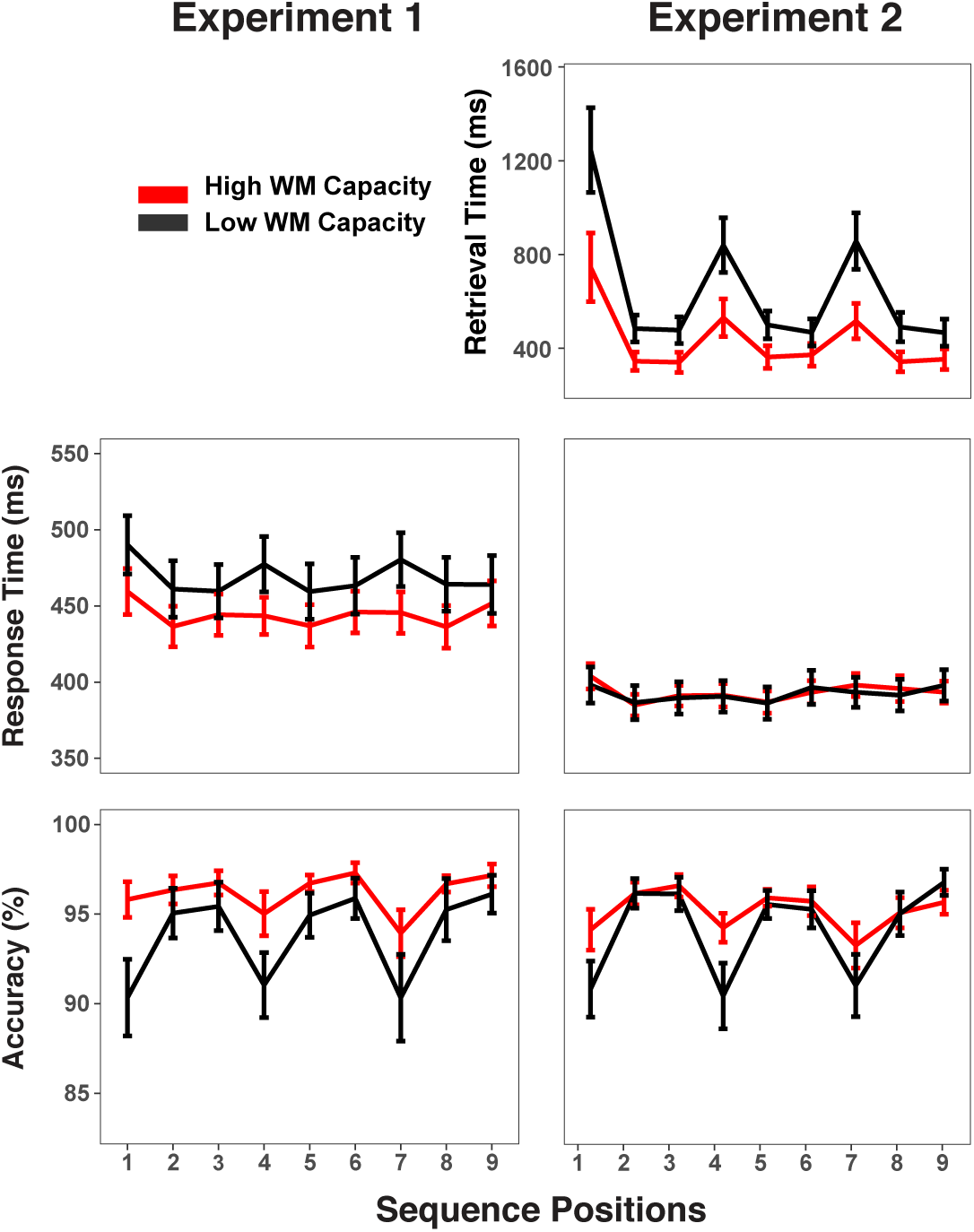
Experiments 1 and 2 behavioral results. RTs and accuracy show marked sequence structure effects in Experiment 1. In Experiment 2, a substantial portion of the RT sequence structure effect is shifted into the self-paced retrieval period. Error bars show within-subject 95% confidence intervals.

For Experiment 2, where participants moved to the next trial only after indicating per key-press that they had retrieved the upcoming orientation, the chunking pattern in RTs, while still reliable was with 5 ms (*SD*= 9.3) was even smaller than in Experiment 1 for RTs, *F*(1,28)=8.05, *MSE*=168.79, *p*=.01, but with 3.5% (*SD*=4.9%) equally strong for errors, *F*(1,28)=18.41, *MSE*<0.01, *p*<.001. There was a very strong chunk-boundary effect of 372 ms (*SD*=361) in the self-paced retrieval times, *F*(1,28)=35.43, *MSE*=234826, *p*<.001. Thus, overall the self-paced retrieval instruction was effective in moving a large share of the retrieval demands out of the EEG-recorded preparation period. At the same time, the additional opportunity for self-paced retrieval did not change the fact that participants made more retrieval errors at transition points.

Figure 2 also indicates that individuals with low WM capacity (i.e., determined through median split) have larger chunk-boundary effects than individuals with high WM capacity. To increase statistical power for detecting such individual-differences effects within our design, we combined accuracy scores across both experiments. The WM-group by chunk-boundary interaction was reliable across both experiments, *F*(1,56)=6.63, *MSE* <.01, *p*=.01, and it was close to reliable in each of the individual experiments, Exp. 1: *F*(1,28)=3.24, *MSE*<.01, *p*=.08, Exp. 2: *F*(1,28)=3.40, *MSE*<.01, *p*=.08. Given that the use of self-paced retrieval in Experiment 2 eliminated much of the retrieval demands from RTs, a meaningful test for RT effects can be conducted only in Experiment 1, where the WM-group by chunk-boundary interaction was in the predicted direction, albeit not reliable, *F*(1,28)=1.51, *MSE*<168.79, *p*=.22. For Experiment 2, the WM-group by chunk-boundary interaction was nearly reliable for self-paced retrieval times, *F*(1,28)=4.19, *MSE*=976921, *p*=.05. Thus, overall, there is an across experiments consistent pattern that people with lower WM capacity have greater difficulty at chunk boundaries.

### Basic Elements: Orientations

Individual orientations served as the basic elements of the pre-instructed sequences that had to be retrieved and actively maintained in WM for the comparison with the test probe. Based on previous work, we hypothesized that the spatial pattern of alpha-band (8-12 Hz) oscillations across electrodes contains information about individual orientations (i.e., 45° vs. 90° vs.135° orientation). Indeed, we found across Experiments 1 and 2 that oscillations centered around the alpha-frequency band allowed sustained decoding of orientations, both late in the preparation period and early in the probe period (cluster-forming threshold *p*<0.05, corrected significance level *p*<0.001; Figure 3a). Moreover, during the probe period, higher posterior probability for elements in the alpha band also predicted faster trial-to-trial responses over and above other constructs (i.e., within-chunk positions and chunk identity), *t*(30)=−3.28, *b*=-0.032, *SE*=0.009, *p*<.05 for Experiment 1, and *t*(30)=-3.51, *b*=-0.016, *SE*=0.004, *p*<.01 for Experiment 2. There were no significant predictive effects during the preparation period for elements, *t*(30)=-0.27, *b*=-0.031 for Experiment 1, and *t*(30)=0.73, *b*=0.01 for Experiment 2. These results are consistent with the previous reports that specific WM representations can be decoded from scalp-distributed alpha activity during the maintenance period. It is noteworthy that a unique aspect of the current results is that we find alpha-encoded representations of stimuli retrieved from long-term memory rather than concurrently or recently encoded from the environment. In addition, during the early probe period, the current element was also decoded from theta-band (4-7 Hz) activity. This result is consistent with previous findings in which stimulus-evoked theta oscillations encoded spatial properties of visually presented stimulus (Foster et al., 2016).

**Figure 3.**
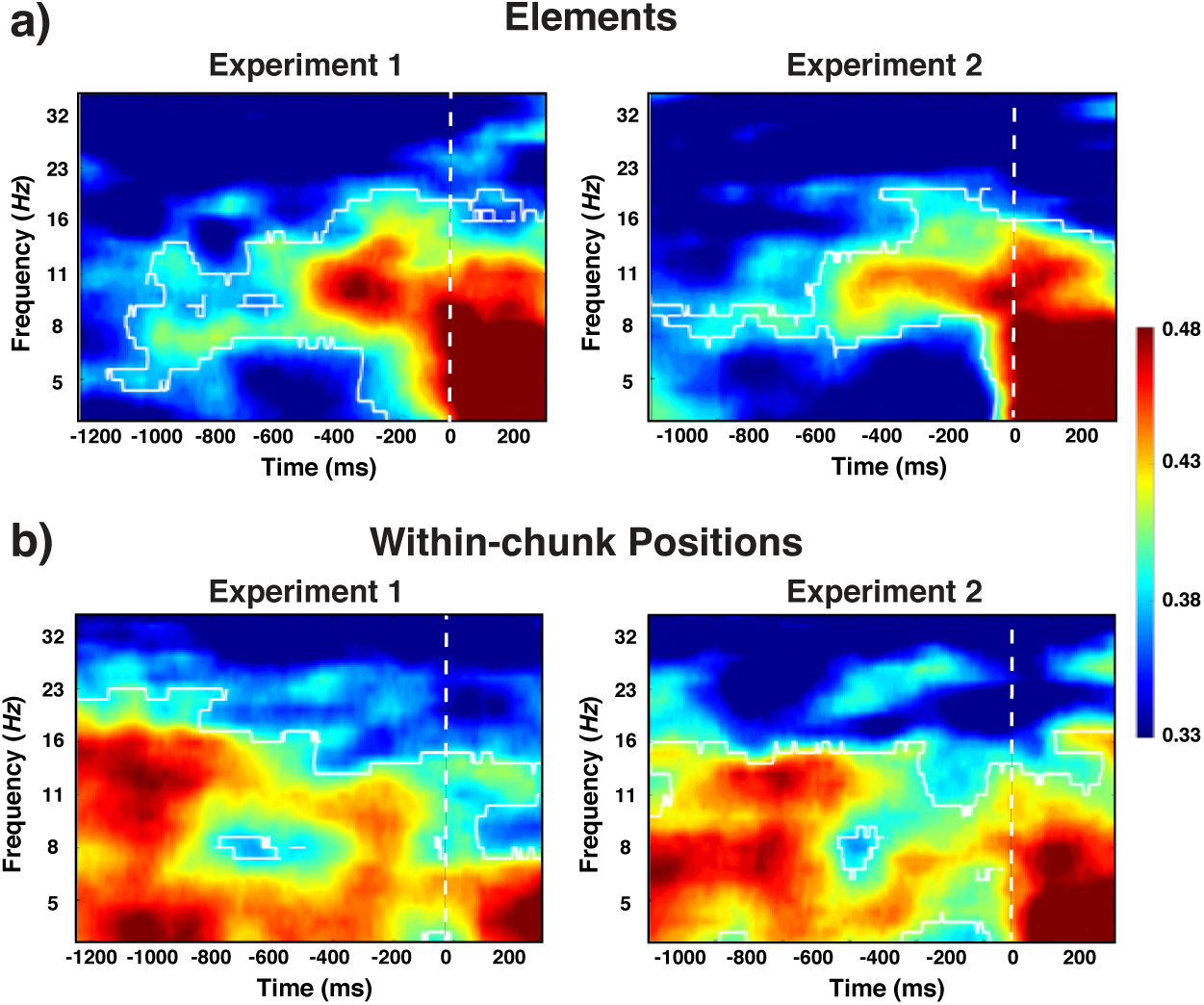
Experiment 1 and 2 decoding results as a function of frequency and time for elements/orientations (a) and within-chunk positions (b). The legend shows the decoding accuracy in probability. Chance level is *p*=.33. The regions enclosed by white lines show significant decoding accuracy after cluster correction.

### Decoding Within-chunk positions

One of our central questions is to what degree we can find evidence for content-independent representations of the current within-chunk position. Based on the previous literature, we hypothesized that theta-band (4-7 Hz) oscillations are a prime candidate as a carrier of such information (Hsieh et al., 2011). Indeed, across Experiments 1 and 2, our decoding results showed that within-chunk position information was present in the broader band of oscillations (4-16 Hz), but with a particular emphasis on the theta-band, from onset of preparation period and stayed elevated during preparation and probe period (cluster-forming threshold *p*<0.05, corrected significance level *p*<0.001; Figure 3b). In the theta-band, robust decoding accuracy was observed at the beginning of the preparation and probe period, whereas, decoding accuracy gradually decreased in the alpha-band as the probe period approached. In addition, more robust evidence for within-chunk position information in the theta-band during the probe period predicted faster RTs in Experiment 1, which used the fixed-paced procedure, *t*(30)=-2.44, *b*=-0.011, *SE*=0.0047, *p*<.05. However, information in the alpha-band was not predictive of RTs, *t*(30)=0.31, *b*=-0.002, *SE*=0.0068, *p*=.76. The theta-band effect was in the same direction but not reliable in Experiment 2, which used the self-paced procedure, *t*(30)=-1.12, *b*=-0.005, *SE*=0.0044. It is possible that the overall reduced variability of RTs (see Figure 2) in the self-paced study may have rendered it less sensitive to capturing critical trial-to-trial variability in performance.

### Ruling Out Processing-Demand Confounds

As mentioned, one important question is to what degree processing-demand confounds across within-chunk positions (e.g., retrieval demands) may produce position-like decoding results. Retrieval demands and supposedly also other processing demands should be considerably reduced in the self-paced procedure used in Experiment 2, relative to the fixed-paced procedure in Experiment 1. Thus, the fact that the decoding results were very similar across the two experiments speaks against the role of such confounds. Nevertheless, to further probe the possible contribution of such effects we can also examine the degree of representational similarity across positions (Grootswagers, Wardle, & Carlson, 2017; Kriegeskorte, Mur, & Bandettini, 2008). Representational similarity analyses (RSA) reveal the nature of decoded information based on the assumption that similar representations recruit analogous neural resources. If the decoding results reflect abstract position codes one would expect that the pattern of EEG activity is equally distinct across all three positions (i.e., equally dissimilar to each other). In contrast, if heightened retrieval/processing demands during position 1 are responsible for the decoding results then we would expect that position 1 is particularly distinct from positions 2 and 3, whereas these two positions should be more similar to each other.

As evident in Figure 4, the pattern of classification probabilities across all positions suggests that position 1 is particularly distinct from positions 2 and 3 during the preparation period. However, during the probe period the three positions appeared to be equally dissimilar to each other. As a statistical test of these observations, we used hierarchical, linear regression models to predict classification probabilities simultaneously with both the unique-position-1 model matrix and the discrete-position model matrix (see bottom panels in Figure 4). During the preparation period, both models significantly predicted the observed confusion matrix, unique-position-1: *t*(30)=5.36; discrete: *t*(30)=3.38; for Experiment 1, unique-position-1: *t*(30)=5.56; discrete: *t*(30)=3.53; for Experiment 2. However, only the discrete model significantly predicted the observed pattern during the probe period, unique-position-1: *t*(30)=0.08; discrete: *t*(30)=2.59 for Experiment 1; unique-position-1: *t*(30)=0.02; discrete: *t*(30)=4.23, for Experiment 2.

**Figure 4.**
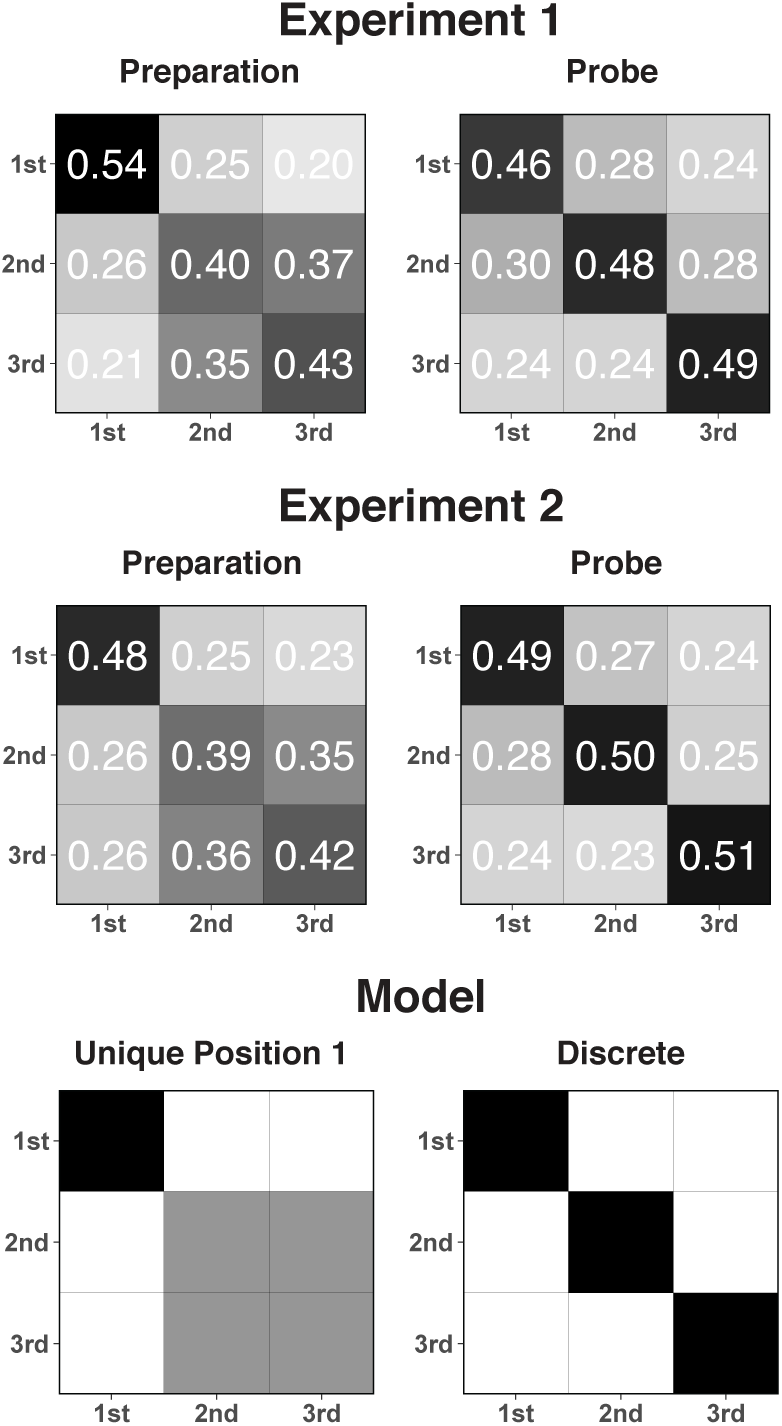
Confusion matrixes for position decoding results for Experiments 1 and 2 and separately for the preparation and the probe period. The bottom panel shows the model matrices used to test the unique-position-1 account against the discrete-position-coding account (see text for details).

As a final check for potential processing-demand effects, we also examined the average theta power at mid-frontal electrodes (Fz and Cz electrodes). Theta power has been repeatedly shown to reflect cognitive control demands and therefore should be sensitive to a position/control demand confound (Cavanagh & Frank, 2014; M. X. Cohen & Donner, 2013). During both the preparation and probe period, the average theta power was not significantly modulated between the first and the remaining positions, *t*(30)=1.00, *b*= 0.49, SE=0.49, and *t*(30)=0.17, *b*=0.12, SE=0.70 for Experiment 1; *t*(30)=0.59, *b*=0.17, SE=0.29 and *t*(30)=-0.16, *b*=-0.07, SE=0.45 for Experiment 2. Thus, these results indicate that the observed decoding results are unlikely to be driven by a confound between positions and control/retrieval demands. Also, they rule out the possibility that position decoding in the theta band simply reflects univariate and consistent changes across individuals that are solely driven by frontal theta power modulation. Rather, the topography of theta oscillations appears to be unique to each position within a chunk.

To summarize, for the preparation period, the pattern of decoding results is somewhat ambiguous with regard to the question whether they truly reflect characteristics of the sequential representation or position-correlated processing-demands. It is worth noting though that a unique-position-1 pattern by itself is not necessarily a strong indicator of such a confound. People may use less precise position representations during preparation (i.e., differentiating mainly between the beginning and later portions of a chunk) and arrive at a more fine-grained representation only during the probe period, when the actual response is imminent. The fact that we obtained similar results across Experiments 1 and 2, despite the very different processing demands during the preparation periods across these experiments, makes it more likely that these are true representational effects. Also, the fact that frontal theta power was modulated by within-chunk position neither during the preparation period, nor during the probe period speaks against the effect of processing demand confounds. Finally, additional evidence that is inconsistent with a processing-demand interpretation of the position-decoding effects will be provided when we analyze effects of working-memory differences on decoding results.

Irrespective of the interpretation of results during the preparation period, the probe-period results suggest that theta-activity captures actual position codes. The clear evidence of position coding during the probe period, but not the preparation period, is consistent with recent behavioral results indicating that people tend to update their current position within a larger sequence when they actually execute an individual, sequential element, not when they prepare for it (Mayr, Kleffner-Canucci, Kikumoto, & Redford, 2014).

### Decoding of Chunk Identity and Position

In addition to the element-level representations, we examined to what degree information about Level 2 representations can be extracted from the EEG signal. As sequences were constructed from three different chunks, each of which occurred at three different positions within the different larger sequences, we looked for information about chunk identity and chunk position. Indeed, we found that the identity of chunks was coded during the preparation period (cluster-forming threshold *p*<0.05, corrected significance level *p*<.05; Figure 5). Across both Experiments 1 and 2, the effect was most sustained following 600 ms prior to the onset of the test probe in the alpha-band (8-12 Hz). In Experiment 2, there was also above-chance decoding in the theta band around the time of stimulus representation; an effect that we will not further interpret given that it was not found in Experiment 1. Generally, only very little chunk-identity information was detectable during the probe period.

**Figure 5.**
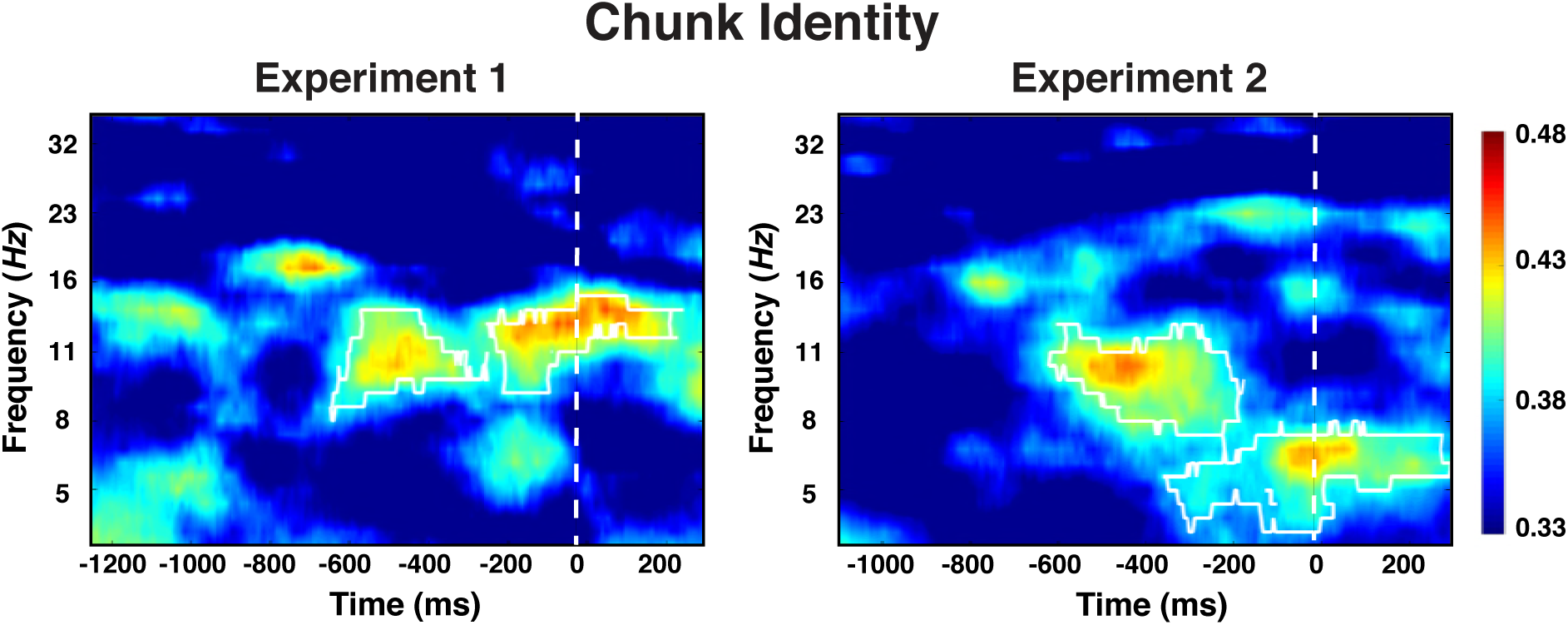
Experiment 1 and 2 decoding results as a function of frequency and time for chunk identity. The legend shows the decoding accuracy in probability. Chance level is *p*=.33. The regions enclosed by white lines show significant decoding accuracy after cluster correction.

One important theoretical question is to what degree chunk representations are active mainly during chunk transitions (i.e., during the first position in each chunk) or are required equally across all positions, potentially to constrain lower-level representations. To address this question, we examined to what degree the chunk-specific neural patterns recur across all three within-chunk positions. Therefore, we summarized the decoding results for each position separately with classifiers trained on the data with all positions. As evident in the Figure 6, while there is some variability in the strength of expression of chunk information for positions 2 and 3 across the two experiments, there is no evidence that chunk information is present only for position 1. A contrast between positions 1 versus positions 2 and 3 produced a reliable effect in neither experiment, Experiment 1: *t*=.02, *p*=.99, Experiment 2: *t*(30)=. 88, *p*=.38. The contrast between positions 2 and 3 did produce a reliable difference in Experiment 1, *t*(30)=2.17, *p*<.05, but not in Experiment 2, *t* (30)=-1.12, *p*=.27. Thus overall, information about the current chunk is represented not only when a new chunk needs to be accessed, but at least to some degree also while within-chunk elements are being processed.

**Figure 6.**
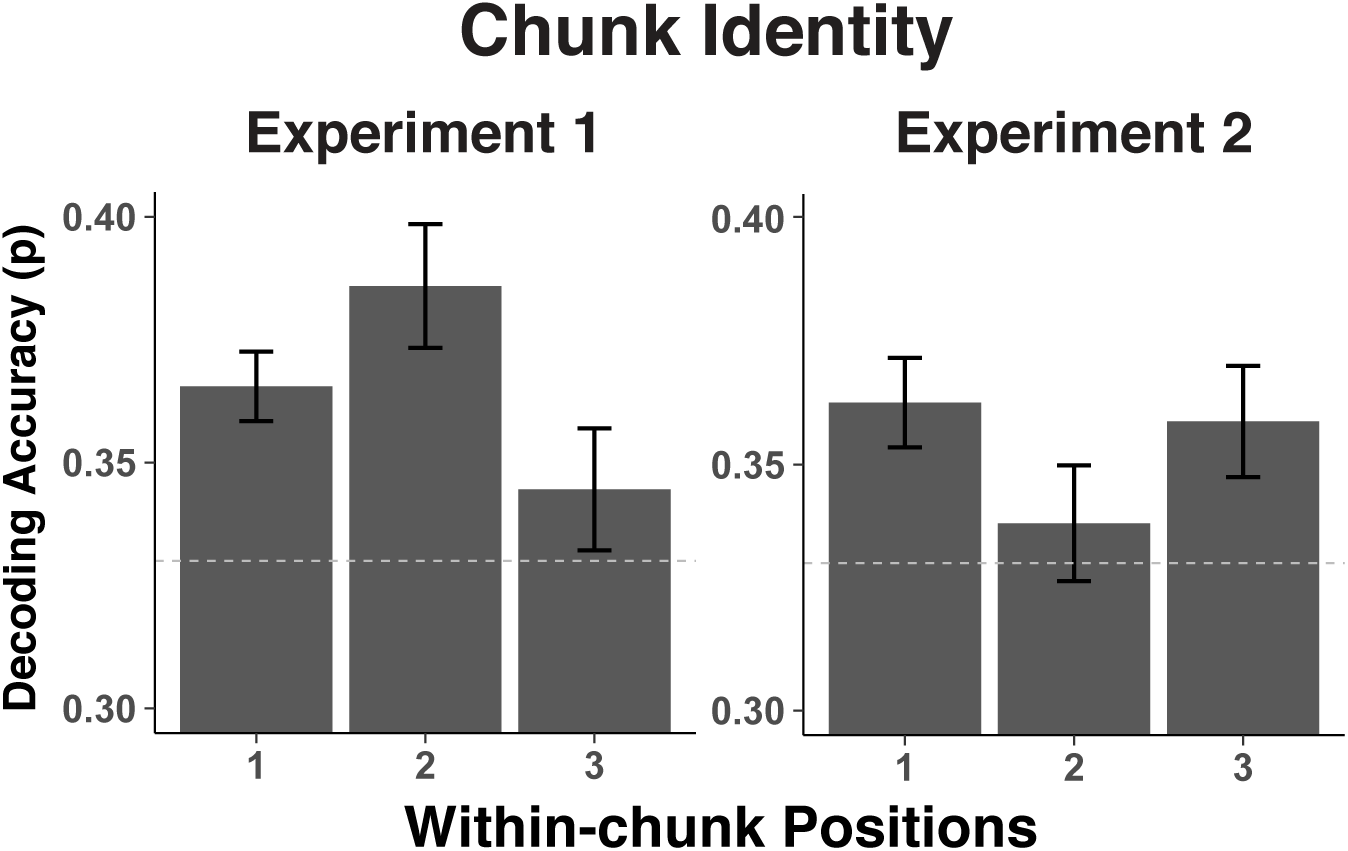
Experiment 1 and 2 decoding accuracy for chunk identity across within-chunk positions. The error bars reflect within-subject 95% confidence intervals.

We also attempted to decode information about the ordinal position of each chunk within the larger sequence. As we have done for the preceding decoding analyses, we initially averaged across all other aspects, including within-chunk positions. In this analysis, we found no evidence of reliable decoding of chunk position codes. However, it is possible that chunk order information is activated only during chunk transitions (i.e., when the next chunk needs to be activated). Such an activation pattern would be diluted when averaging across within-chunk positions. Indeed, when we attempted to decode chunk positions separately for each within-chunk position, we obtained significant clusters in the alpha-band immediately after transitions (i.e., during the preparation period of position 1) and, in the theta-band towards the end of the probe period of position 3, that is just prior to the transition (cluster-forming threshold *p*<0.05, corrected significance level, *p*<0.01 for both alpha and theta effects in both studies; see Figure 7). No significant clusters were identified at the second position of chunks. As these results were not a-priori predicted they need to be considered with care. However, they are consistent across the two experiments and they are consistent with the plausible assumption that chunk position representations are recruited as a new chunk needs to be established, likely as cue towards the retrieval of the upcoming chunk’s identity.

**Figure 7.**
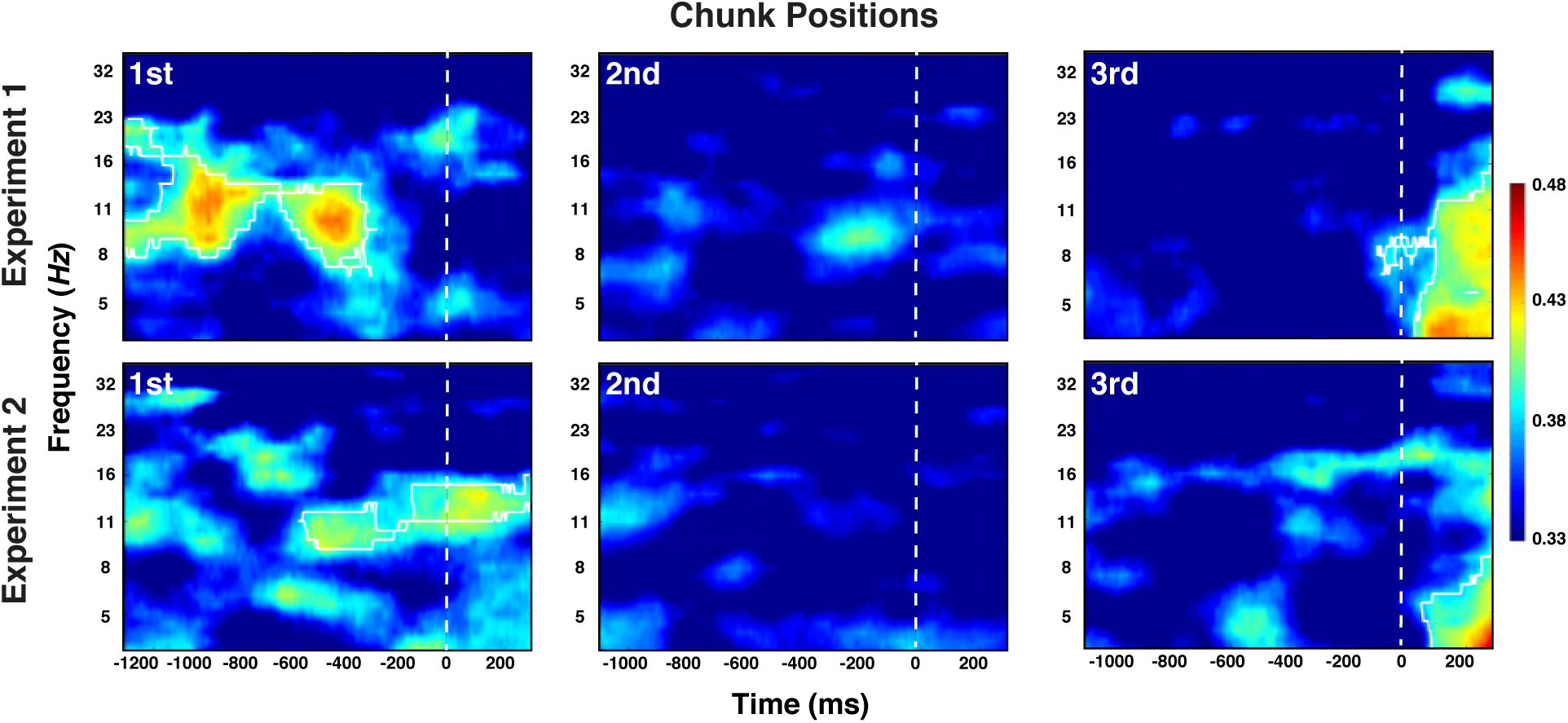
Experiment 1 and 2 decoding results as a function of frequency and time for chunk position, shown separately for the within-chunk positions 1, 2, and 3. The legend shows the decoding accuracy in probability. Chance level is *p*=.33. The regions enclosed by white lines show significant decoding accuracy after cluster correction.

### Individual difference and WM capacity

When combining Experiments 1 and 2 our sample size is—for EEG research—reasonably large and therefore allows us to probe the role of individual differences in the decoding of serial-order control codes. Behavioral results had indicated larger chunk-boundary costs in RTs, errors, or retrieval times for low-WM individuals than for high-WM individuals (see Figure 2). This pattern is consistent with the hypothesis that WM selectively constrains the representation of level-2 codes of chunk identity and/or position. To further test this hypothesis, we examined the degree to which decoding accuracy, as an indicator of representational strength, differed as a function of WM capacity for each of the different serial-order control codes. For Level-1 representations, we focused here on the a-priori predicted frequency bands (alpha band for elements and theta band for positions). For Level-2 representations, we focused on those bands and periods (i.e., preparation vs. probe) for which we had found consistent above-chance decoding in the general analyses. As apparent from Table 1, neither in Experiment 1 nor in Experiment 2 did the representational strength for Level-1, basic element and position codes differ between individuals with high versus low WM capacity. For Level-2 chunk identity codes however, decoding accuracy was robustly modulated by individuals’ WM capacity during the preparation period, which was the only period during which chunk identity information was detectable. As apparent in Figure 8, in both experiments, there was no above-chance chunk-identity information in low-WM participants, but robust information for high-WM participants.

**Figure 8.**
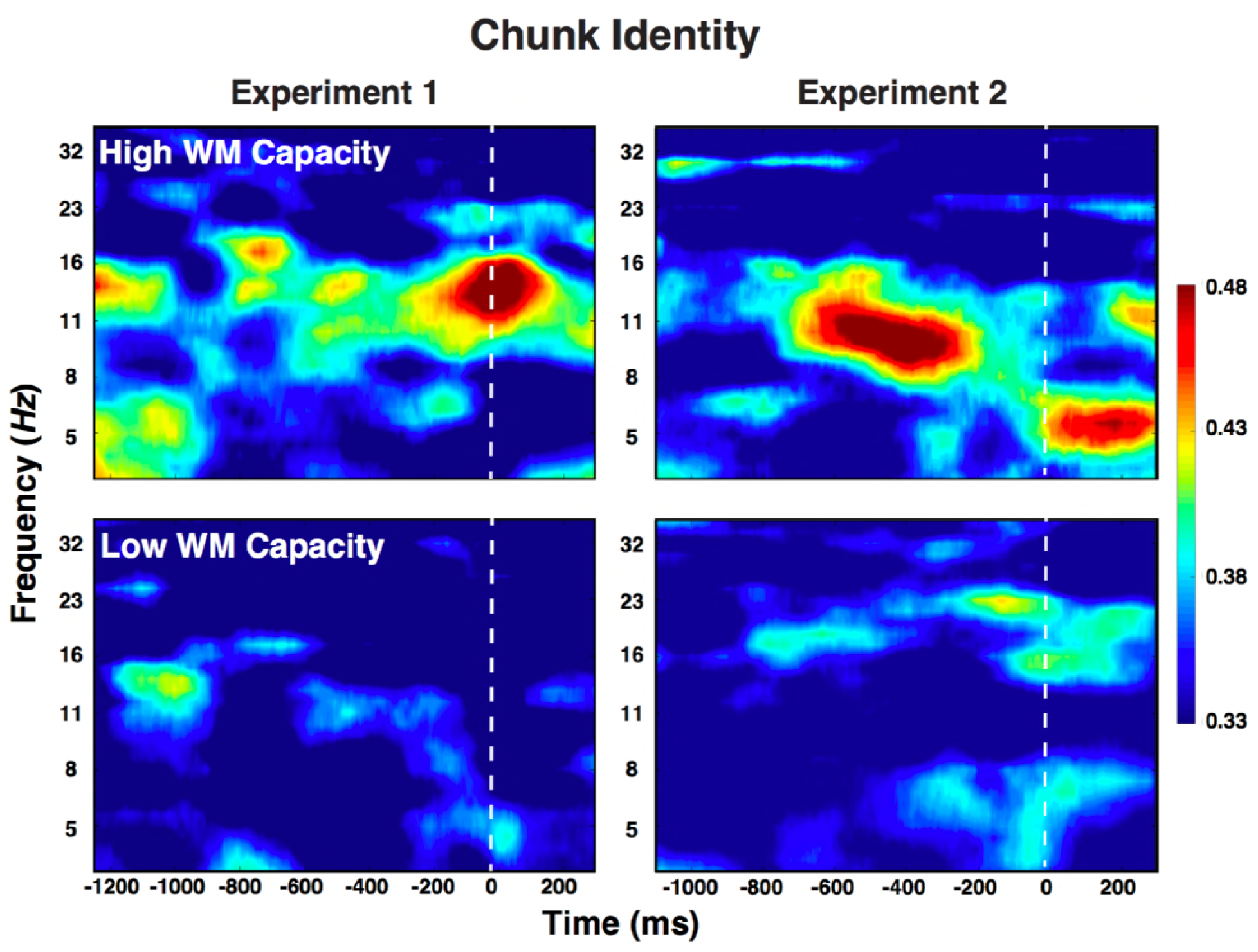
Experiment 1 and 2 decoding results as a function of frequency and time for chunk identity for individuals with low versus high WM capacity.

**Figure 9.**
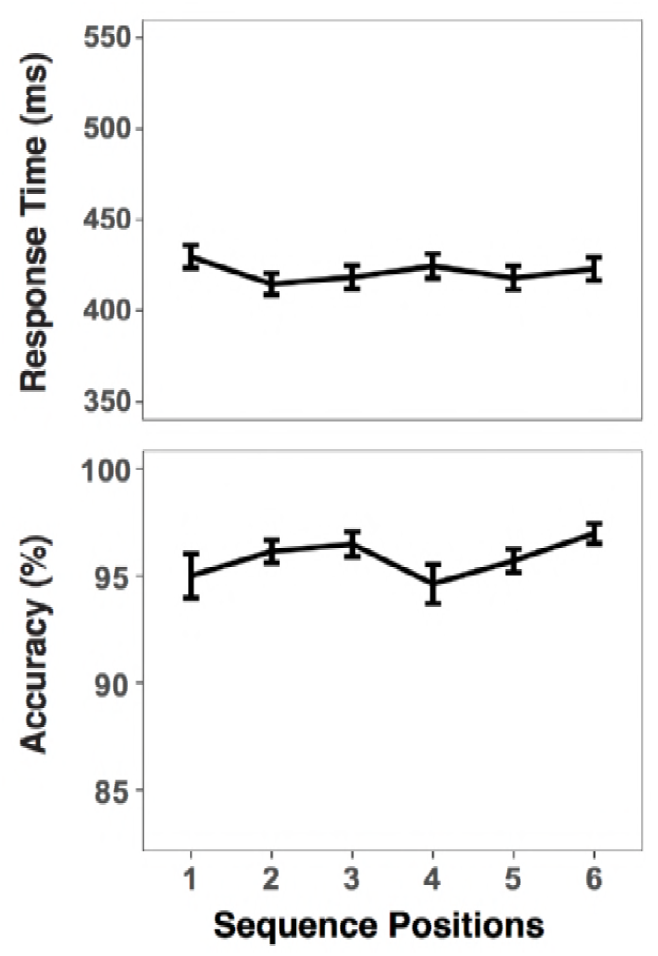
RTs and accuracies as a function of sequence positions. Error bars show within-subject 95% confidence intervals.

**Table 1.** Decoding accuracy for each feature in individuals with high versus low working-memory capacity.

|  |  | Experiment 1 |  |  |  |
| --- | --- | --- | --- | --- | --- |
| Level 1 |  | High WM | Low WM | *t* | *p* |
| Element | Prep | 0.425(0.03) | 0.412(0.02) | 0.329 | 0.744 |
|  | Probe | 0.427(0.04) | 0.463(0.02) | -0.816 | 0.421 |
| Position | Prep | 0.443(0.03) | 0.388(0.03) | 1.408 | 0.170 |
|  | Probe | 0.477(0.02) | 0.411(0.03) | 1.530 | 0.137 |
| Level 2 |  |  |  |  |  |
| Chunk | Prep | **0.380(0.02)** | **0.325(0.01)** | **2.503** | **0.018** |
|  | Probe | 0.368(0.03) | 0.323(0.04) | 0.910 | 0.370 |
| Position | Pos. 3 Probe | 0.422(0.04) | 0.357(0.04) | 1.196 | 0.241 |
|  | Pos. 1 Prep. | 0.360(0.02) | 0.371(0.02) | 0.380 | 0.707 |
|  |  | Experiment 2 |  |  |  |
| Level 1 |  | High WM | Low WM | *t* | *p* |
| Element | Prep | 0.390(0.02) | 0.407(0.03) | -0.403 | 0.689 |
|  | Probe | 0.445(0.04) | 0.435(0.03) | 0.207 | 0.837 |
| Position | Prep | 0.422(0.02) | 0.433(0.02) | -0.380 | 0.701 |
|  | Probe | 0.468(0.04) | 0.479(0.02) | -0.231 | 0.818 |
| Level 2 |  |  |  |  |  |
| Chunk | Prep | **0.407(0.01)** | **0.326(0.02)** | **3.427** | **0.002** |
|  | Probe | 0.373(0.03) | 0.337(0.02) | 1.045 | 0.305 |
| Position | Pos. 3 Probe | 0.384(0.03) | 0.377(0.03) | 0.162 | 0.872 |
|  | Pos. 1 Prep. | 0.348(0.02) | 0.374(0.03) | -0.321 | 0.750 |

*Note.* For each feature, decoding accuracy was averaged for either the preparation or the probe period. For Level-2 position codes, decoding accuracy was compared for the position-3 probe period (early in the transition) and the position-1 preparation period (late in the transition, see Figure 7).

In order to provide a test of the differential effect of WM on the decoding of Level-2 chunk identity, but not Level-1 element identity, we also included decoding results across both experiments into an ANOVA with experiment and level as between-subject factors and level (i.e., chunk vs. element) as within-subject factor. The critical interaction between level and WM group was reliable, *F*(1,58)=4.30, *p*=.044, *eta*^2^=.07, whereas none of the interactions with the experiment factor reached significance, all *F*s<1.

For chunk position codes, we focused on the periods at chunk beginnings (alpha band) and endings (theta band) during which reliable position decoding had been detected across the entire group. However, we found no consistent WM group differences here (see Table 1). Thus, overall, the pattern of WM effects on decoding accuracy was not compatible with the hypothesis that WM affects serial-order control representations in a uniform manner. Rather, WM capacity seems to selectively constrain the ability to represent the current chunk context while preparing for the next, within-chunk element.

Finally, the pattern of individual differences in decoding accuracy allows us to revisit the concern that the decoding of within-chunk positions may be driven by processing-demand confounds. The behavioral results had indicated that low-WM individuals had greater difficulty than high-WM individuals at the first chunk position (Figure 2), suggesting that any position-related processing-demand effects were particularly strong for that group. Thus, if position decoding is driven by differential processing demands then we would expect stronger decoding accuracy for low than for high WM individuals. However, for both Experiments 1 and 2, within-chunk position decoding accuracy was equally robust (see Table 1), adding to the earlier presented arguments (see *Ruling out Processing Demand Confounds*) that within-chunk decoding reflects the sequential representation rather than position-specific processing demands.

### Experiment 3

A potential qualification of our position decoding results from Experiments 1 and 2 is that within-chunk sequences contained element-to-element transitions that were invariant across chunks, which in principle could allow a simple, associative chaining process to contribute to sequential representations. Although it is not obvious that chaining of elements can produce decoding patterns consistent with ordinal positions, it would nevertheless be reassuring to generalize our results to sequences that do not allow associative chaining. In Experiment 3, we therefore used sequences constructed from a set of six different chunks, such that for each sequence, element-to-element transitions were ambiguous and could not be learned via associative chaining (A. Cohen, Ivry, & Keele, 1990).

The behavioral results (Figure 10) show that effects of chunk-boundary effects on RTs were with 10.73 ms (SD=8.5) small, albeit reliable, *F*(1,19)=19.48, *MSE*=969.88, *p*<.001. Also, for errors the chunk boundary effect was with 1.12 % (*SD*=3.13) considerably smaller than in the preceding experiments, albeit still reliable, *F*(1,19)=5.33, *MSE*<.01, *p*<.001. Different from the self-pacing procedure in Experiment 2, we achieved the relatively small boundary effects by allowing repeatable pre-practice with each sequence. Thus, this experiment provides yet another opportunity to examine representations of ordinal positions, while minimizing potential processing-load confounds.

**Figure 10.**
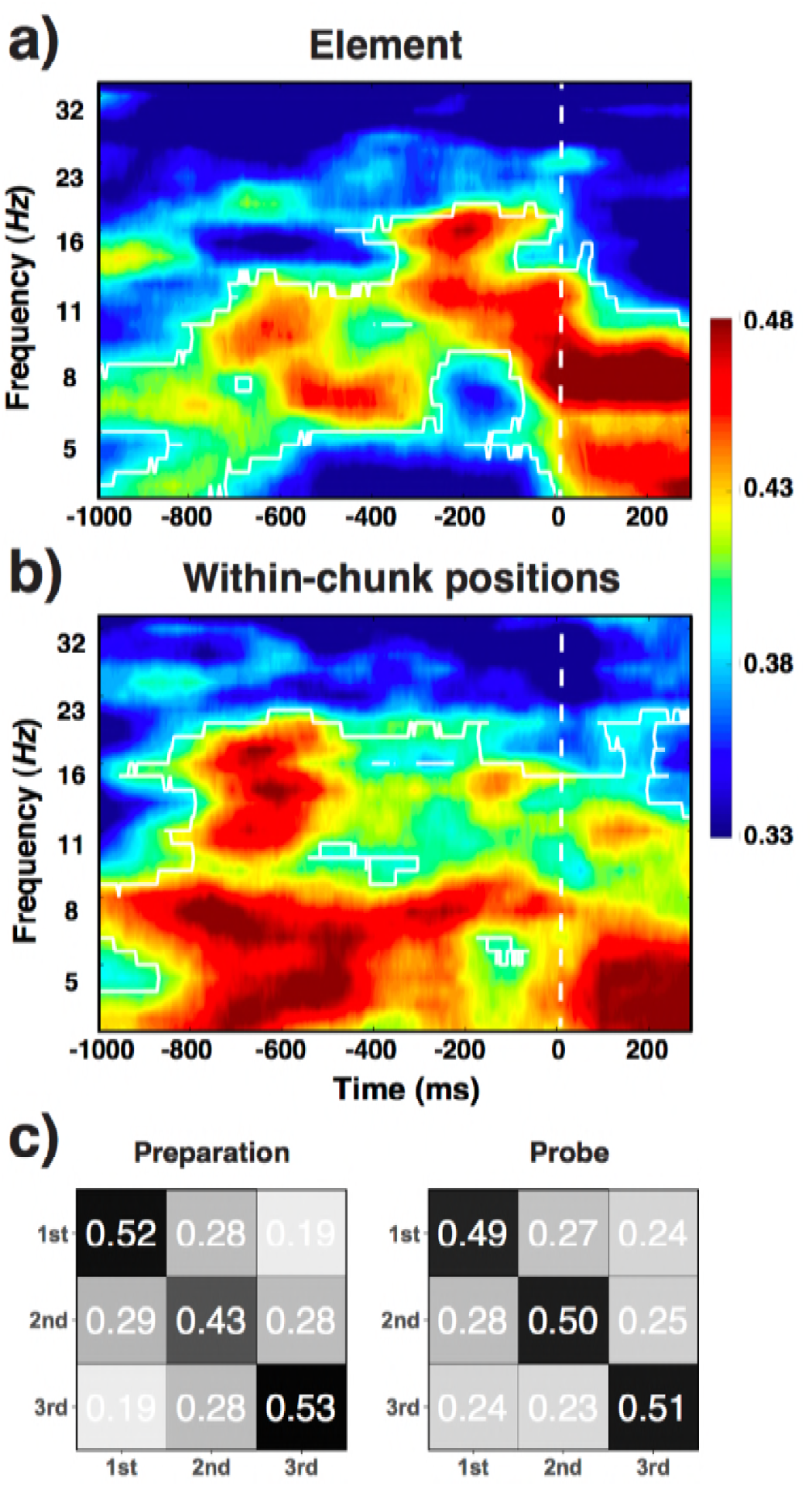
a) Experiment 3 decoding results as a function of frequency and time for element/orientation identity and within-chunk position. The legend shows the decoding accuracy in probability. Chance level is *p*=.33. The regions enclosed by white lines show significant decoding accuracy after cluster correction. b) Confusion matrixes for position decoding results from Experiment 3, separately for the preparation and the probe period.

The decoding results fully replicated the findings for Level-1 codes from Experiments 1 and 2 (Figure 10a). Alpha-band activity during the preparation period and the probe period contained robust information about the specific identity of basic elements (i.e., orientations; cluster-forming threshold *p*<0.05, corrected significance level *p*<0.01). Ordinal positions were also successfully decoded in the theta band both for the preparation and the probe period, and to a lesser degree also in the alpha band during the preparation period (cluster-forming threshold *p*<0.05, corrected significance level *p*<0.01).

As in the previous experiments, we performed RSA analyses to evaluate the possible influence of retrieval confounds on the decoding of positions (Figure 10b). The pattern of theta activity was distinct across all three within-chunk positions, unique-position-1: *t*(20)=3.02; discrete: *t*(20)=1.75). However, we found such a pattern also for the preparation period, for which we had observed a greater degree of confusion between the first position and the remaining positions in Experiments 1 and 2, unique-position-1: *t*(20)=3.30; discrete: *t*(20)=1.90). The additional practice with each sequence, or the need for more robust sequential representations in the face of greater between-chunk interference, may have induced participants to engage in a greater degree of preparation in Experiment 3. Combined, these results indicate that reliable position decoding is not limited to “chainable” sequences (as used in Experiments 1 and 2) and they provide further evidence that position decoding cannot be explained in terms of processing/retrieval demand confounds.

As in Experiments 1 (and numerically in Experiment 2) we found higher decoding accuracy of within-chunk position codes in theta-band predicted faster RTs during the preparation period instead of probe period, *t*(30)=-2.09, *b*=-0.009, *SE*=0.0047, and *t*(30)=-0.51, *B* =-0.003, *SE*=0.006. For element codes in the alpha-band, we replicated the same pattern of results from Experiment 1 and 2, *t*(30)=0.27, *b*=0.002, *SE*=0.007 for the preparation period, and *t*(30)=-1.64, *b*=-0.005, *SE*=0.0044 for the probe period.

## Discussion

According to a longstanding theoretical tradition, sequential behavior is organized in terms of hierarchically structured representations that are independent of the actual sequence content (Lashley, 1951). In principle however, what appears to be hierarchically organized behavior can also be produced through associative chaining, where the sequential knowledge resides in associations between successive elements or complex, “hidden-layer” integrations of such elements (Botvinick & Plaut, 2004). Chaining-based representations require neither content-independent codes nor an explicit, hierarchical structure.

Our results present strong evidence that explicitly instructed sequential action is in fact controlled via hierarchically organized, abstract codes. Evidence for sequence-extraneous control codes was particularly convincing for representations of the positions of basic sequential elements (Level 1 position codes). In addition, we found direct evidence for neural signals that represent the identity of chunks and potentially even signals that coded the position of chunks within the larger sequence. Our results complement findings from both animal neurophysiological work and human fMRI studies that have provided initial evidence for abstract position codes and for their neuroanatomical substrate (Averbeck & Lee, 2007; Berdyyeva & Olson, 2010; Desrochers et al., 2015; Fujii & Graybiel, 2003; Kalm & Norris, 2014; Ninokura, Mushiake, & Tanji, 2003). The combination of EEG decoding and an explicit sequencing paradigm allowed us to track control codes on different hierarchical levels in a time-resolved manner, while avoiding important confounds that pose validity challenges to position-code results in earlier fMRI work (Kalm & Norris, 2017a, 2017b).

### Position and Element Codes

Based on previous evidence, we had expected that basic sequential elements (i.e., different orientations) would be captured predominantly in alpha-band activity (Foster et al., 2016; Fukuda et al., 2016; Palva, Monto, Kulashekhar, & Palva, 2010; Sauseng et al., 2009); Foster et al, 2016; Fukuda et al, 2016), whereas within-chunk position codes would be represented in theta-band activity (Heusser et al., 2016; Hsieh et al., 2011; Hsieh, Gruber, Jenkins, & Ranganath, 2014).

As expected, information about the basic elements (orientations) was indeed expressed strongest in the alpha band during the preparation period, whereas information about the within-chunk positions of basic elements was expressed strongest in the theta band, both in the preparation period and the probe period. These results are the first to show within humans, that theta activity not only responds to serial-memory load (Hsieh et al., 2011), but actually contains abstract, content-independent position information. In particular, detailed analyses of the representational similarity structure among the position codes provided clear evidence for equidistant, ordinal position codes during the probe period. However, in Experiments 1 and 2, for the preparation period, the difference between position 1 versus positions 2 and 3 was particularly dominant. As elaborated in the Results section, we have good reasons to assume that this reflects the nature of the representation used during preparation. In addition, we found evidence for ordinal position coding for both during preparation and during the probe period in Experiment 3, where additional practice may have produced more robust chunk representations.

The finding that ordinal position information is encoded in the theta band is consistent with Lashley’s original idea that serial order can be established through content-independent, position codes. However, in general there was more a quantitative than a strict qualitative dissociation between the representation of the identity of basic elements and serial-order information. For example, across all three experiments we found in the alpha band an unexpected, short phase of position decoding around the beginning of the preparation period. The temporal signature of the decoded information at the beginning of the preparation period may indicate that it reflects the effort of activating the position code from LTM. Instead, the more sustained, theta-based activity likely represents the established position context. Conversely, during the probe period, we observed a strong presence of element information in the theta band—in the absence of any alpha-band element decoding. Indeed, the decoding of low-level spatial features of visually presented items has been previously found in stimulus-evoked theta-band oscillations (e.g., Foster et al., 2016). Thus, this activity may reflect the fact that bottom-triggered representations interact with the current sequential context.

### Chunk-level Representations

Larger sequences can only be represented through small sets of position codes when they are broken down into manageable chunks. These chunks in turn, need to be represented and organized in terms of their serial order. Our results provide initial evidence that just as for within-chunk information, there were distinct representations for identity and serial order at the chunk level. While chunk identity information was represented in the alpha band, there was some indication that order information was carried also by theta band activity. This pattern may suggest a more general dissociation between the representation of *what* needs to be done (elements or chunks) and *when* it needs to be done (positions of elements or chunks). It also indicates recursivity in terms of the principles by which information is represented across levels (Fitch & Martins, 2014).

There is a considerable body of fMRI imaging work that tries to map out the neuroanatomy of different levels of the cognitive control architecture (Badre, 2008; Farooqui et al., 2012; Koechlin & Summerfield, 2007). While our EEG-based results provide no information about localization of levels, it does provide information about the temporal flow of control within a hierarchical structure. One important question in this regard is to what degree higher-level representations (i.e., chunks and chunk positions) and low-level representations (i.e., elements and element positions) are activated sequentially, such that the first are needed to initiate the latter. Alternatively, higher-level and lower-level representations could be activated in parallel, potentially constraining each other (Ranti et al., 2015). Our results indicate that information about the position of a chunk within the larger sequence is activated only at chunk transition points, presumably as a cue to retrieve the next chunk identity. In contrast, information about chunks themselves was activated somewhat in parallel to information about the next orientation and its ordinal position for within-chunk element. This latter result is more consistent with the parallel-activation account.

### Working Memory Constraints

Individual differences in WM capacity predict peoples’ ability to perform complex, sequential tasks (e.g., Carpenter et al., 1990). Yet, there is little research that directly examines which aspects of serial-order control are affected by WM constraints. In principle, such constraints might affect all involved representations in a uniform manner. Alternatively, constraints might be specific to particular representations. For example, according to one theoretical conceptualization (Oberauer, 2009), WM provides a protected space for binding a limited number of elements to a structural frame (e.g., objects to spatial locations or sequential positions). Assuming that WM is used to tie basic elements to position codes, one would expect little effect of WM capacity on these representations as long as the size of chunks does not exceed WM capacity. In fact, across two experiments we found that individual differences in WM capacity were not systematically related to Level-1 element or position codes. However, hierarchical sequences also require that WM maintains the larger context, such as of the current chunk or chunk position within the overall sequence. On the behavioral level, we found that individual differences in WM were particularly strong for between-chunk transitions, suggesting that individuals with less WM capacity had a harder time “finding” the next chunk. The EEG results suggested that this may be due to the fact that only high-capacity individuals were able to execute within-chunk elements, while maintaining a robust representation of the current chunk identity. Thus, at least for short sub-sequences, WM constraints are specific to Level-2 chunk representations, which in turn are critical to maintain one’s position within a larger sequential context. Further, these results also indicate that the answer to the question to what degree Level-2 representations are active in parallel to Level-1 representations comes with some additional nuance. The exact architecture of control seems to depend on available WM capacity, where only high-capacity individuals exhibit strong evidence for parallel Level-1 and Level-2 activity.

### Qualifications

The current results support the existence of abstract, serial-order control codes. They do not rule out that associative chaining representations govern behavior in different types of sequencing situations, such as after extensive practice of a particular sequence, or when position information is not useful (Botvinick & Plaut, 2004; Davachi & DuBrow, 2015; Keele, Ivry, Mayr, Hazeltine, & Heuer, 2003). Strictly speaking, we cannot even rule out chaining-type representations in our Experiments 1 and 2, where the makeup of our sequences would have allowed such representations to emerge. However, in Experiment 3 we eliminated any utility of associative links. The fact that our position-decoding results were essentially identical across these experiments, indicates that even if associative-chaining representations did emerge in Experiments 1 and 2, they did not affect the relevance of position-code representations.

Schapiro, Kustner, and Turk-Browne (2012) suggested as a test for associative chaining to compare the forward and backward predictability between successive representations early and late in practice. Forward predictability is given when element *N* is more similar to the post-learning pattern of element *N*+1 compared to the extent to which the pattern elicited by *N*+1 is similar to the pre-learning pattern of N. We carried out tests of forward predictability separately for elements at positions 1 to 2 and 2 to 3 in each of our sequences, examining the first half versus the second half of each block. In no case did we find any indication of a forward predictability pattern (all *p*> 0.29). Even though these results rest on accepting the null hypothesis they are at least consistent with the conclusion that chaining was not a major factor.

Another qualification is that our procedures were particularly geared towards a robust decoding of within-chunk elements and position codes by presenting each element and each position combination across the three chunks in each sequence. Had we assessed each chunk in separate blocks, decoding would have been affected by common temporal context, likely making it more difficult to extract pure position-specific or element-specific information. While the use of three chunks per sequence also allowed us to decode Level-2 representations, the design was less geared towards the robust assessment of chunk identify or order information. In particular, the results regarding chunk position codes need to be considered with some caution. The fact that chunk position decoding was limited to the boundaries between chunks (i.e., for chunks at the end of third position and the beginning of first position) is theoretically plausible, but also makes it particularly difficult to rule out retrieval or other processing-demand confounds.

Finally, an obvious limitation of our work is that while the EEG-based decoding approach yields high temporal resolution, it provides no information about the neuroanatomical origin of the observed rhythmic activity. In this regard, previous research suggests some hypotheses. In serial memory tasks, a frontally distributed enhancement of theta power has been observed that is commonly associated with the medial and lateral PFC (Hsieh et al., 2011; Jensen & Lisman, 2005; Jensen & Tesche, 2002). This finding converges with monkey electrophysiology (Averbeck & Lee, 2007; Berdyyeva & Olson, 2010) and human fMRI studies (Desrochers et al., 2015; Kalm & Norris, 2017b; Ninokura et al., 2003) in which neural activity in the PFC showed sensitivity to both serial positions and hierarchical control demands (Badre & D’Esposito, 2009). Furthermore, several studies suggest that the hippocampus, where theta oscillations are prominent, is a critical structure for coding temporal context (Heusser et al., 2016; Hyman, Zilli, Paley, & Hasselmo, 2005; Jones & Wilson, 2005; Siapas, Lubenov, & Wilson, 2005). More specifically, it is possible that the hippocampus drives prefrontal theta modulation (Ferino, Thierry, & Glowinski, 1987).

### Conclusion

Our results suggest that time-resolved decoding analyses with EEG can be utilized to characterize abstract control representations. Specifically, we provide strong evidence that oscillatory activity in the theta band contains information about sequential position codes that is independent of the representation of the content of a sequence. In addition, we provide initial evidence about the representation of chunk-level information, as well as about how WM constraints affect representations of chunk identity.

## Method

### Participants

A total of 88 students of the University of Oregon participated in this research and received compensation of $10 per hour. Four participants from Experiment 1, two participants from Experiment 2, and two participants from Experiment 3 were excluded because of a higher than 25% trial rejection rate due to EEG artifacts. The final samples contained 30 participants each for Experiments 1 and 2, and 20 participants for Experiment 3. All experimental protocols were approved by the University of Oregon’s Human Subjects Committee. Past work has reported robust EEG decoding of perceptual or working memory representations with sample sizes of less than 20 subjects (e.g., Foster et al., 2016). Given that this is the first study to look at the decoding of abstract control representations, we tried to establish the robustness of our key findings by both using larger samples for two of our experiments (*N*=30 in Experiments 1 and 2) and by replicating key results across experiments. The larger sample sizes and consistent sequence designs across Experiments 1 and 2 also allowed us to explore individual differences in decoding accuracy by combining data across these experiments.

### Task and Stimuli

On each individual trial of the experiment, participants were asked to compare an orientation associated with the current position within a larger sequence with an orientation probe stimulus, presented on the screen (Figure 1b). These orientations were presented as lines within a circle (2°in diameter) in which lines were tilted in 45°, 90°, or 135° angles. Each trial contained a preparation and a probe period. During the preparation period, participants were encouraged to pre-retrieve the corresponding element for the upcoming test. This phase was indicated by the fixation cross (0.25° in diameter) presented at the center of the screen. The preparation period was set to be 1250ms, 1100ms, or 1000ms for Experiments 1, 2, and 3 respectively. During the probe period, a potential orientation stimulus was presented, and participants indicated per key press whether or not the probe matched the element corresponding to the current position within the sequence (“z” key pressed with the left index finger for non-matches; “? /” key with the right-index finger for matches). In order to encourage preparation, mild time-pressure was induced by requiring responses within a 700ms time window; later responses were counted as errors. After the response followed a jittered, inter-trial-interval (ITI) of 500 to 900ms. Additionally, for Experiment 2, the preparation period was preceded by a self-paced “retrieval period”, where participants initiated the preparation period by pressing the space bar in a self-paced manner, once they had retrieved the next element. When participants made an error, the assigned sequence was displayed, highlighting the element corresponding to the current position. This error feedback was presented until the correct response was executed and it remained on the screen into the following trial. Participants were encouraged to make judgments to the probe as fast and accurate as possible. All task-related visual stimuli were generated in Matlab (Mathworks) using the Psychophysics Toolbox (Brainard, 1997) and were presented on a 17-inch CRT monitor (refresh rate: 60 Hz) at a viewing distance of 100 cm.

### Sequences and Blocks

Prior to each experimental block of trials, a new sequence was instructed. The instruction screen displayed the entire sequence in the form of a horizontal sequence with small spaces between independent chunks. In Experiments 1 and 2, participants were given a new 9-element sequence made of three unique chunks of three unique elements (ABC, BCA, CAB) for each block of trials (Figure 1a). Counterbalancing the order of all chunks yields six unique sequences, each of which was presented equally often within 24 experimental blocks of 54 trials each (i.e., six sequence cycles per block). For Experiment 3, participants were instructed six-element sequence made of two, 3-element chunks, selected from a total of six possible chunks (ABC, ACB, CAB, CBA, BAC, BCA). In this experiment, sequences were constructed such that elements never repeated across chunk boundaries, which yields a total of 12 possible sequences. Participants performed 36 experimental blocks of 42 trials each (i.e., 7 sequence cycles per block). Note, that in all three experiments, the procedures ensured that each element (orientation), each position, chunk identity, and chunk position are combined equally often across blocks.

Before the actual experiment begins, participants were given practice blocks. In Experiments 1 and 2, each of three chunks were tested in the aforementioned match/mismatch task, and participants performed 3 practice blocks of 27 trials each (i.e., 4 chunk cycles per block). In Experiment 3, we attempted to provide more robust pretraining with individual chunks with the hope to counteract position-dependent processing demands (e.g., Kalm & Norris, 2017a). To this end, a first chunk was randomly selected from the pool of six possible chunks and repeatedly tested until it was reproduced with 100% accuracy by clicking on the three possible orientations in the correct order for a given chunk. At this point, the next chunk was added and tested, randomly intermixed with the first chunk until the 100% performance criterion was achieved. This procedure was repeated until the performance for all chunks reached the performance criterion. Given that each error within a practice block led to the repetition of that block, this procedure ensured considerable exposure to the set of chunks. Its duration was approximately 20 minutes for the majority of participants (compared to under 5 minutes for Experiments 1 and 2).

### Measurement of WM capacity

To assess participants’ WM capacity, we chose a standard, visual change-detection paradigm (Luck & Vogel, 1997) because it imposes no serial-order demands. Participants were presented arrays of four or eight colored squares (0.65°×0.65°) at random locations during a 100 ms encoding interval. After a delay period of 900 ms, a single probe at the location of one of the squares of the encoding set was presented. Participants had to indicate whether or not the color of the probe matched the color of the original square. There were 160 trials each for set-sizes 4 and 8. The capacity was computed as K=S × (H - F), where K is the memory capacity, S is the size of the array, H is the observed hit rate and F is the false alarm rate (Cowan, 2001).

### EEG recordings and preprocessing

Similar as in Kikumoto and Mayr (2017) Electroencephalographic (EEG) activity was recorded from 20 tin electrodes held in place by an elastic cap (Electrocap International) using the International 10/20 system. The 10/20 sites F3, Fz, F4, T3, C3, CZ, C4, T4, P3, PZ, P4, T5, T6, O1, and O2 were used along with five nonstandard sites: OL midway between T5 and O1; OR midway between T6 and O2; PO3 midway between P3 and OL; PO4 midway between P4 and OR; and POz midway between PO3 and PO4. The left-mastoid was used as reference for all recording sites. Data were re-referenced off-line to the average of all scalp electrodes. Electrodes placed ∼1cm to the left and right of the external canthi of each eye recorded horizontal electrooculogram (EOG) to measure horizontal saccades. To detect blinks, vertical EOG was recorded from an electrode placed beneath the left eye and reference to the left mastoid. The EEG and EOG were amplified with an SA Instrumentation amplifier with a bandpass of 0.01–80 *Hz* and were digitized at 250 *Hz* in LabView 6.1 running on a PC. We used the Signal Processing, EEGLAB (Delorme & Makeig, 2004) toolboxes, and custom-made codes for EEG processing in MATLAB. Trials including blinks (>80uv, window size=200ms, window step=50ms), large eye movements (>1°, window size=200ms, window step=10ms), and blocking of signals (range=-0.05 *u*v to 0.05uv, window size=200ms) were excluded from further analysis, resulting in an average of 920 trials per participants.

### EEG analysis

Single-trial EEG data were decomposed into their time-frequency representation by a complex wavelet convolution. Following Mike X Cohen (2014), the power spectrum of the EEG signal was obtained through fast Fourier transformation of the raw EEG signal. The power spectrum was then convolved with a series of complex Morlet wavelets (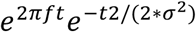), where *t* is time, *f* is frequency increased from 4 to 35 Hz in 32 logarithmically spaced steps, and σ defines the width of each frequency band, set according to *n*/*2πf,* where *n* increased from 3 to 10. The incremental number of wavelet cycles was used to balance temporal and frequency precisions as the function of frequency of the wavelet and the logarithmic scaling was used to keep the width across frequency bands approximately constant. After this is done in the frequency-domain, we took the inverse of the Fourier transform, resulting in complex signals in the time-domain. A frequency band-specific estimate was defined as the squared magnitude of the convolved signal *Z*(real([*z*(*t*)]^2^ + imag[*z*(*t*)]2) for instantaneous power. For frequency-band specific analyses, these signals were averaged within pre-selected ranges of frequency values (4-7 Hz for theta-band, 8-12 Hz for alpha-band).

### Multivariate pattern analysis

In order to investigate whether the pattern of oscillatory power carries information about the representations of hierarchical sequence, we performed a series of linear discriminant analyses. First, all artifact-free trials, excluding errors and post-error trials were randomly partitioned into four blocks. To prevent the inflation of decoding accuracy, the number of observations for each to-be-decoded condition (e.g., first vs. second vs. third position within a chunk) was equated within and across blocks. Next, the average oscillatory power was computed for each electrode with 100 ms (i.e., 25 time samples) sliding window. To remove the effects of potential processes that affect multiple electrodes in a uniform manner (e.g., sensory adaptations, (Kalm & Norris, 2017a), all data points were transformed into z-score. This resulted in an 1 × b × e × s matrix of power values, where l is the number of conditions, b is the number of blocks, e is the number of electrodes, and s is the number of time samples. The decoding analysis was performed for each frequency value (or frequency-band) independently over time. Each block was assigned as a training set, which was used to develop a linear discriminant function to decode the condition label of an independent test set. We used a hold-one-block-out cross-validation procedure, in which power estimates from three of four randomly partitioned blocks served as the training set and those from the remaining block served as the test set. This process was repeated until each block had served as the test set. To improve the signal-to-noise ratio of the decoding analysis, we introduced an iterative procedure in which the entire process described above was repeated 30 times to obtain the average across iterations. In addition to the overall classification accuracy, estimates of the posterior probability for the correct label was retained for each individual trial.

### Statistical Analyses

Each decoding analysis was performed independently within participants. However, to account for multiple comparisons, we conducted non-parametric permutation tests when evaluating decoding results in the time-frequency space on the level of the group (Maris & Oostenveld, 2007). First, clusters of samples adjacent across the temporal and frequency dimension in which the decoding accuracy was higher than chance level (i.e., *t*-value > 2.0) were identified. Then, empirical cluster-level statistics were obtained by taking the sum of *t*-values in the largest cluster. Finally, nonparametric statistical tests were performed by calculating a cluster *p*-value under the permutation distribution of cluster-level statistics, which were obtained by Monte Carlo iterations of decoding analyses with randomly shuffled condition labels. Both the cluster entry threshold and the cluster significance threshold were set to *p*<.05, one-sided. A priori hypotheses about the difference between WM memory group was assessed by two sample t-tests after averaging the decoding accuracy within frequency band. The reported effect sizes corresponded to the mean effect size within subject ± SEM across participants, and the p values were obtained from the second-level analyses obtained across participants.

To investigate how the trial-to-trial variability in the quality of control codes relates to sequencing performance, we used multilevel modeling to predict single trial RTs by sequence representations. First, RTs were log-transformed and residualized by the linear and quadratic trends of experimental trials and blocks as well as the effect of within-chunk ordinal positions with two orthogonal contrast codes (see *Behavior* section) and the congruence of test probes. Then, we tested models predicting RTs with single trial posterior probabilities from the decoding analysis averaged within the hypothesized frequency bands for each construct. We ran separate models for the preparation period and the probe period with trials nested within participants. All models estimated both random intercepts and slopes of all constructs simultaneously.

## Acknowledgements

This work was supported by NSF grant 1734264 awarded to Ulrich Mayr.

